# Reconstitution of a Biofilm Adhesin System from a Sulfate-Reducing Bacterium in *Pseudomonas fluorescens*

**DOI:** 10.1101/2023.11.22.568322

**Authors:** Amruta A. Karbelkar, Maria E. Font, T. Jarrod Smith, Holger Sondermann, George A. O’Toole

## Abstract

Biofilms of the sulfate reducing bacterium (SRB) *Desulfovibrio vulgaris* Hildenborough (DvH) can facilitate metal corrosion in various industrial and environmental settings leading to substantial economic losses; however, the mechanisms of biofilm formation by DvH are not yet well-understood. Evidence suggests that a large adhesin, DvhA, may be contributing to biofilm formation in DvH. The *dvhA* gene and its neighbors encode proteins that resemble the Lap system, which regulates biofilm formation by *Pseudomonas fluorescens*, including a LapG-like protease DvhG and effector protein DvhD, which has key differences from the previously described LapD. By expressing the Lap-like adhesion components of DvH in *P. fluorescens*, our data support the model that the N-terminal fragment of the large adhesin DvhA serves as an adhesin “retention module” and is the target of the DvhG/DvhD regulatory module, thereby controlling cell-surface location of the adhesin. By heterologously expressing the DvhG/DvhD-like proteins in a *P. fluorescens* background lacking native regulation (Δ*lapG*Δ*lapD*) we also show that cell surface regulation of the adhesin is dependent upon the intracellular levels of c-di-GMP. This study provides insight into the key players responsible for biofilm formation by DvH, thereby expanding our understanding of Lap-like systems.

**Significance:** Corrosion leads to ∼2.5 trillion US dollars in economic losses, 20% of which are estimated to be microbially induced. Biofilms of sulfate reducing bacteria (SRB), especially the genus *Desulfovibrio*, are important members of the corrosion consortium and accelerate deterioration of metals. Understanding how biofilms are formed by SRB can provide important clues to mitigate this challenge. In this study, we used genetic and biochemical tools to investigate the mechanism of biofilm formation by *Desulfovibrio vulgaris* Hildenborough. Our study reveals critical genes responsible for regulating the secretion of a large adhesin known to be required for biofilm formation.

## Introduction

Bacteria predominantly exist as biofilms in various environmental niches (1). Free-swimming bacteria can sense diverse inputs and respond to those cues by transitioning to surface-attached biofilms (2). Initial steps in biofilm formation require various factors, including motility, pili and protein adhesins, and exopolysaccharides (2). Biofilms eventually become embedded in an adhesive matrix consisting of polysaccharides, proteins, and extracellular DNA (3). While the triggers and mechanisms of biofilm formation are varied among different bacteria, these processes are ultimately governed by bacterial nucleotide secondary messenger c-di-GMP in many microbes (4).

Biofilms of sulfate-reducing bacteria (SRB), particularly of the genus *Desulfovibrio*, have been shown to play an important role in bioremediation of heavy metals (5) and sulfur cycling in various ecological settings (6). These microbes can accept electrons from extracellular electron donors, such as minerals or electrodes (7). As a result, their biofilms can be used as catalysts in bioelectrochemical systems for wastewater treatment (8) and energy production (9). SRB are also responsible for substantial economic losses as their biofilms can clog gas and oil pipelines and cause microbially induced corrosion of metal structures (10). Therefore, understanding SRB biofilms and how they form has major implications for both natural and engineered systems.

The Gram-negative, obligate anaerobe, *Desulfovibrio vulgaris* Hildenborough has been used as a model organism to understand the mechanisms and effects of biofilm formation by SRB (11). For example, quorum sensing affects various metabolic processes in *D. vulgaris* including an increase in biofilm formation (12). Several studies have also investigated the components of biofilm matrix. For example, analysis of gene expression levels in *D. vulgaris* biofilms showed up to a 1.8-fold increase in exopolysaccharide biosynthesis genes compared to planktonic cells (13). The biofilm matrix of *D. vulgaris* has been shown to be protein-rich (observed as filaments by electron microscopy) and low in carbohydrates (14). Transcriptomic and proteomic analysis of *D. vulgaris* biofilms revealed two large proteins, DVU1012 and DVU1545, in the biofilm matrix, both of which contribute to biofilm formation (15). Isolation and identification of membrane-associated proteins implicated DVU1012 as being localized to the outer membrane in two separate studies (16, 17). More recently, a study focusing on a *D. vulgaris* variant that unexpectedly lost its ability to form a biofilm discovered that a spontaneous mutation in an ABC transporter component DVU1017 was responsible for this phenotype. Incidentally, the DVU1017 protein is situated several genes downstream of the gene coding for the large protein DVU1012 (18). This study strongly suggested that DVU1017 is part of a non-canonical type I secretion system (T1SS) (19), called the Lap system, that has been thoroughly characterized for its role in biofilm formation in the environmental microbe *Pseudomonas fluorescens* Pf0-1. That is, DVU1012 was proposed to be the LapA-like protein of *D. vulgaris* (18).

Biofilm formation by *P. fluorescens* Pf0-1 is mediated primarily by a large adhesive protein, LapA which is localized to the outer membrane via anchoring in the LapE outer membrane protein (**Figure 1a**) (20). The Lap system is comprised of an ABC-transporter LapB, fusion protein LapC, and outer membrane porin LapE that allows LapA to traverse from the cytoplasm to the outer membrane via its C-terminal secretion domain. Our previous biochemical studies and comparison to the N-terminus of another LapA-like protein (21) indicates that the N-terminus of LapA can fold into a retention module (RM) which allows LapA to remain anchored on the OM (22). LapA displayed on the outer membrane allows cells to adhere to a surface and promote biofilm formation. Whether LapA remains tethered to the OM depends upon two regulatory proteins: a periplasmic protease LapG and an effector protein LapD, the latter of which can bind c-di-GMP (23, 24). The LapG protease can cleave LapA downstream of the retention module leading to a loss of the adhesin from the cell surface. Under conditions of high intracellular c-di-GMP, this dinucleotide binds to the cytoplasmic domain of LapD, which in turn leads to sequestration of LapG in a LapD-LapG complex in the periplasm. As a result, LapA remains tethered to the OM. Under low c-di-GMP conditions, c-di-GMP is not available for LapD binding by its cytoplasmic domain, and this receptor undergoes a conformational change allowing the release of LapG in the periplasm (**Figure 1a**). LapG then cleaves LapA, causing release of LapA and a loss of the biofilm (23, 25).

**Figure 1:**
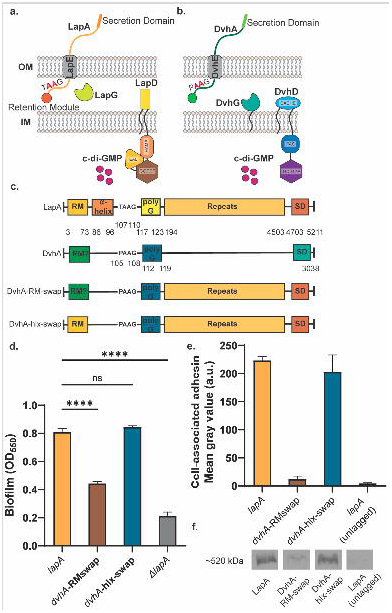
Simplified model of two-step Type 1 secretion system (T1SS). a) T1SS in *Pseudomonas fluorescens* Pf0-1. The large adhesin LapA is localized to the outer membrane (OM) via a retention module, allowing the bacterium to form a biofilm. The localization of LapA to the OM is regulated by a periplasmic protease LapG and a cyclic di-GMP (cdG) effector protein LapD. At high intracellular cdG levels, the catalytically inactive EAL domain of LapD binds cdG leading to a conformational change whereby the LapD periplasmic domain sequesters LapG, thereby retaining LapA on the OM. When c-di-GMP levels are low, LapD is in its autoinhibited conformation, which releases LapG to cleave LapA at dialanine motif (shown in red), resulting in the release of LapA to the extracellular environment and loss of biofilm formation. b) Proposed model of T1SS in *Desulfovibrio vulgaris* Hildenborough. DvhA is a LapA-like adhesin localized to the OM. DvhG is a LapG-like protease that can cleave DvhA at the dialanine site (shown in red), however, unlike the periplasmic-localized LapG, DvhG is hypothesized to be inner membrane (IM) bound. DvhD is a structurally different but functionally analogous LapD-like protein containing a catalytically inactive HD-GYP domain that can bind c-di-GMP and thus regulate DvhA localization and biofilm formation in *D. vulgaris* Hildenborough. c) Simplified protein architecture representing different domains and the dialanine site of Pf0-1 adhesin LapA, DvH adhesin DvhA and fusion proteins DvhA-RM-swap and DvhA-hlx-swap (RM - retention module; polyG – polyglycine linker; SD – C-terminal secretion domain). Protein domains are not to scale. d) Biofilm formed in K10T medium at 24 hours by strains expressing the adhesins DvhA-RM-swap and DvhA-hlx-swap compared to the native adhesin LapA in Pf0-1, all in the Δ*lapG*Δ*lapD* background strain. e) Quantification of cell surface levels of HA-tagged adhesins at 24 hours in K10T medium. f) Western blots indicating total cellular levels of the different adhesins. Statistical analysis was performed using one way ANOVA multiple comparisons (ns, p > 0.05; *, p ≤ 0.05; **, p ≤ 0.01; ***, p ≤ 0.001; ****, p ≤ 0.0001); ****, P<0.0001).

In this study, we build on a bioinformatic analysis that revealed an unusual Lap-like system in *D. vulgaris* Hildenborough. We investigate the mechanism of biofilm formation mediated by the Lap-like system of *D. vulgaris* via heterologous expression to introduce components of the SRB Lap system into *P. fluorescens* Pf0-1. This analysis provides new insight into a new group of Lap-like adhesion systems.

## Results

### Bioinformatic analyses identify a Lap-like system in SRB and other organisms

To date, our knowledge of how Lap-like biofilm adhesins function arises from work done largely in fluorescent pseudomonads, with some additional studies in *Bordetella bronchiseptica*, *Legionella pneumophila* and *Vibrio cholerae* (26–29). To broaden our understanding of other Lap systems, we performed a bioinformatics search comparing genomes of bacteria that encode LapG- and LapD-like proteins, which are canonical Lap proteins of *P. fluorescens* (**Fig. 1a**). The Simple Modular Architecture Research Tool (SMART) was used to identify proteins with a C93 Peptidase domain (LapG) or LapD_MoxY_N domain (LapD). Most bacteria investigated using this method encode both LapG- and LapD-like components (**Fig S1a**), but several genera, such as sulfur reducing bacteria of *Desulfovibrio*, only encode LapG-like proteins (**Fig S1b**). Manual inspection of several SRB genomes, including *D. vulgaris* Hildenborough revealed genetic architecture similar to the Lap system of *P. fluorescens* (**Fig S1c**).

The genomic environment around the genes coding for LapG-like proteins in *D. vulgaris* predicted genes encoding a T1SS, a LapG-like protease (DVU1019, DvhG), a LapB-like ABC transporter (DVU1017, DvhB), a LapC-like periplasmic adaptor (DVU1018, DvhC), a LapE-like OM porin (DVU1013, DvhE) and a LapA-like large adhesin (DVU1012, DvhA) (18) (**Figure S1c**). Interestingly, while we did not find a LapD-like protein based on sequence similarity, the *D. vulgaris* genome encodes the DVU1020 protein, that we predict may function similarly to LapD of *P. fluourescens* Pf0-1 (abbreviated Pf0-1) and that we call DvhD.

### Proposed model of biofilm formation by the Lap system of *D. vulgaris* Hildenborough

While we were performing the bioinformatic analysis outlined above, work from De León et al. reported functional data supporting a role for the DvHABCDG system in biofilm formation (18). Specifically, a spontaneous point mutation arose in the *dvhB* gene (coding for a component of the T1SS) rendering the strain biofilm-defective with no detectable DvhA on the surface of the cell. Based on our bioinformatic analysis and study of the Pf0-1 Lap system, the work from De León et al. supported a model for biofilm formation via the large adhesin protein DvhA by *D. vulgaris*.

In this model (**Figure 1b**), we hypothesize that DvhA can be translocated from the cytoplasm to the OM via the components of the T1SS: DvhB, DvhC and DvhE. The N-terminus of the DvhA sequence has spaced glycine residues that suggest possible folding into a retention module as observed for LapA (**Figure S1d,e**). DvhA also possesses an N-terminal dialanine motif, 106AA107, characteristic of LapA-like adhesins and essential for LapG proteolysis in Pf0-1 (30). We hypothesized that DvhG cleaves DvhA at this dialanine site. Interestingly, the DvhG sequence reveals a transmembrane helix which suggests that this protease is inner membrane bound (unlike periplasmic-localized LapG). DvhD is a predicted inner membrane-bound protein which has a periplasmic CACHE domain and a cytoplasmic PAS domain fused with a phosphodiesterase (PDE) ‘HD-GYP’-like domain that, which based on the lack of key residues, is predicted to be catalytically inactive (31). LapD, in contrast, is comprised of a periplasmic PAS domain, a cytoplasmic HAMP domain with catalytically inactive DGC ‘GGDEF’ and PDE ‘EAL’ domain, the latter of which can bind c-di-GMP (**Figure S1f**) (25). While the domains of both proteins are distinct, we predicted that DvhD may be functionally analogous to LapD since they both have periplasmic and cytoplasmic signaling domains that are fused to inactive PDE domains. Thus, we predicted that DvhD can drive signaling through c-di-GMP binding. Overall, our model proposes that DvhG and DvhD can regulate localization of the large adhesin DvhA via c-di-GMP in a manner analogous to LapD/LapG (**Figure 1b**).

To validate the model above, our goal was to establish the heterologous expression of *D. vulgaris* Lap components in Pf0-1 background, with the latter lacking its native LapA machinery. We decided on this approach for the following reasons: 1) While tools for genetic manipulation have been developed for this SRB (32–34), *D. vulgaris* is an obligate anaerobe and is difficult to culture and genetically manipulate compared to Pf0-1, and 2) Our lab has extensively studied the Pf0-1 Lap system, and thus we have a large battery of background strains, cloning tools, and experimental systems that could be deployed to understand the *D. vulgaris* Lap system.

### The N-terminus of DvhA functions as a retention module that promotes biofilm formation

DvhA is a 3038 aa protein that consists of calcium-binding, von Willebrand factor, and glycine-rich domains which are commonly found in LapA-like proteins (**Figure 1c**) (20). At its N-terminus, similar to LapA, we note that DvhA possesses several beta sheets separated by glycine residues – this region could potentially fold into a globular domain (**Figure S1d,e**). Hence, we hypothesized that the DvhA N-terminal sequence folds into a retention module required for biofilm formation. Because of the large size the DvhA protein (∼334kDa), instead of expressing the full-length DvhA in Pf0-1, we constructed two DvhA-LapA protein fusion strains using N-terminal components of DvhA fused to the large C-terminal adhesin domains of LapA; the former would anchor the adhesin to the surface and the latter which would be available to promote surface attachment. Using such constructs, we could assess whether the N-terminal domain of DvhA could anchor the fusion protein to the cell surface to promote biofilm formation and serve as a target for the DvhG protease.

We constructed two different fusion variants. First, we replaced the retention module and LapG proteolysis site of LapA (1M-125S) with the analogous N-terminus of DvhA (1M-119G) and called this fusion protein DvhA-RM-swap (**Figure 1c**). We built this chimeric protein in the Δ*lapG*Δ*lapD* genetic background for two reasons: 1) lack of a functional LapG should result in stoichiometric retention of the fusion proteins on the outer membrane (35), and 2) this strain background would allow us to introduce the DvhG/DvhD proteins to assess their function in the context of the fusion protein.

To test whether the N-terminus of DvhA functions as a retention module, we performed a static biofilm assay with DvhA-RM-swap strain at 24 hours and compared the extent of biofilm formation to the WT Pf0-1 strain expressing LapA as a positive control and no LapA (Δ*lapA*) as a negative control. Our results indicate that DvhA-RM-swap fusion protein can promote biofilm formation (**Fig. 1d**), and although these biofilms were less robust than the WT Pf0-1, they were significantly higher than the negative control (OD_550_ = 0.45 ± 0.008 vs 0.21 ± 0.02, p = 0.0002).

To assess whether the differences in biofilm formation by the strain producing the DvhA-RM-swap variant was due to different amounts of adhesins localized to the cell surface, we performed dot blots using a hemagglutinin (HA) tag that was previously engineered into the chromosome of the gene encoding LapA (36). Cells were grown for 24 hours, centrifuged, washed and OD_600_ normalized before spotting directly on a nitrocellulose membrane. Upon imaging the membrane and quantifying the intensity of the spots, we found that there was significantly less DvhA-RM-swap fusion protein on the cell surface compared to LapA (**Fig. 1e**). This reduced cell-surface DvhA-RM-swap fusion protein was likely due to the fact that the DvhA-RM-swap fusion protein was unstable (Intensity = 1.85 ± 0.36 X10^3^ a.u.; **Fig. 1f**). The amount of surface localized fusion protein was not significantly different from the negative control; however, it tends to be slightly higher (Mean gray value = 18.5 ± 3.97 vs 10.2 ± 1.97, p = 0.098; **Figure 1e**). Together, these data indicate that the N-terminal portion of DvhA can serve to anchor the adhesin to the cell surface to promote biofilm formation, albeit to a lesser degree than the Wt LapA.

Because DvhA-RM-swap did not yield very robust biofilms, we tested another fusion protein, DvhA-hlx-swap (**Figure 1c**). Also constructed in the Pf0-1Δ*lapG*Δ*lapD* background, in this strain, we swapped the residues from LapA that are known to be critical for LapG processing (A81-G123) (30), with sequences from DvhA that we predict are required for DvhG processing (G87-G119), including the di-alanine residues and poly-glycine linker (**Figure 1c**).

To compare the ability of the DvhA-hlx-swap fusion protein to support biofilm formation, we performed a static biofilm assay at 24 hours and compared the extent of biofilm formation by Pf0-1 that expresses the WT LapA in the same Δ*lapG*Δ*lapD* background. Biofilms formed by the strain expressing the DvhA-hlx-swap (OD_550_ = 0.84 ± 0.005) were significantly more robust than DvhA-RM-swap and not significantly different from those formed by Pf0-1 expressing WT LapA (OD_550_ = 0.80 ± 0.01; **Figure 1d**).

Dot blots to probe for surface associated adhesin revealed no difference in signal for the DvhA-hlx-swap fusion and LapA (**Figure 1e**). Western blotting found equivalent signal for LapA and DvhA-hlx-swap fusion (Intensity = 16.3 ± 1.9 X10^3^ a.u. vs 16.5 ± 0.45 X10^3^ a.u.; p > 0.05) in the cytoplasm as well (**Figure 1f**).

Additionally, we confirmed that the DvhA-hlx-swap fusion adhesin uses the T1SS machinery of Pf0-1 since we saw a significantly reduced biofilm and cell surface-associated adhesin upon mutating the ABC transporter, LapB (**Figure S2a, b**). Given the comparable stability of DvhA-hlx-swap to LapA, we decided to explore the mechanics of *D. vulgaris* Lap-like system using the strain Pf0-1 Δ*lapG*Δ*lapD* genetic background expressing the DvhA-hlx-swap variant in all the experiments described below.

### DvhG can cleave the fusion adhesin DvhA-hlx-swap

*P. fluorescens* Pf0-1 LapG is a bacterial transglutaminase-like cysteine protease (BTLCP) that has been shown to cleave LapA adhesin at its 108AA109 dialanine motif (35, 37). This proteolytic activity removes the N-terminal retention module of LapA via a periplasmic cleavage event, causing the remaining protein to “slip” out of the outer membrane LapE channel, thereby leading to a loss of LapA-mediate adhesion. Thus, based on this information, we would predict that DvhG functions similarly to Pf0-1 LapG because 1) DvhG also belongs to the BTLCP family of proteins and (**Figure S3**) 2) the adhesin DvhA possesses a di-alanine site at 106AA107 that could serve as a proteolytic site for DvhG (**Figure S1d**).

First, to test if DvhG can cleave DvhA-hlx-swap, we cloned full length *dvhG* from *D. vulgaris* Hildenborough genome into an arabinose-inducible multicopy plasmid pMQ72 under P_BAD_ promoter (pMQ72-*dvhG*). The P_BAD_ promoter is inducible by arabinose and typically shows some constitutive expression in the absence of induction (38). A histidine tag (6x-His) was also added, resulting in production of a N-terminal His-DvhG protein under control of arabinose induction and that we could detect with α-His antibodies. This plasmid was then transformed into the Pf0-1 Δ*lapG*Δ*lapD dvhA-hlx-swap* strain. Static biofilm assays were performed in the absence or presence of the inducer, 0.2% arabinose, at 24 hours (**Figure 2a**). The results were compared to the same background strain harboring the pMQ72 plasmid, which we will refer to as the empty vector (EV) control, or a strain lacking any plasmid. Our results indicate a significant reduction in the biofilm formed when His-DvhG was expressed under arabinose control compared to the EV control (OD_550_ = 0.89 ± 0.05 vs 0.28 ± 0.006). This phenotype was observed even in the absence of arabinose induction (OD_550_ = 0.31 ± 0.03, p = 0.23 comparing +/- arabinose), indicating leaky expression from this construct. This result suggests that DvhG may be cleaving DvhA-hlx-swap protein causing a loss of the DvhA-hlx-swap fusion protein from the cell surface and a decrease in the biofilm formed.

**Figure 2:**
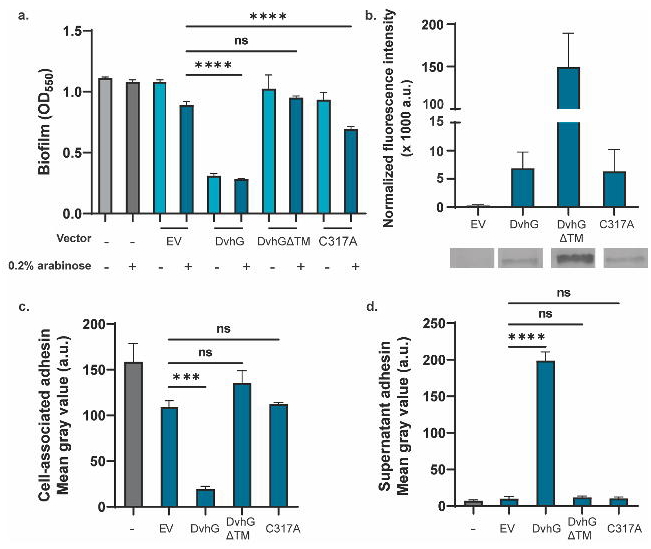
DvhG can cleave DvhA-hlx-swap and diminish biofilm formation. The gene encoding His-tagged DvhG and its variants were cloned in an arabinose inducible promoter and transformed into Pf0-1 Δ*lapG*Δ*lapDdvhA-hlx-swap*. The protease was induced with 0.2% arabinose. a) Biofilm formed in K10T medium at 24 hours in the presence or absence of arabinose with no vector, empty vector (EV), or a plasmid expressing full length DvhG, DvhG lacking the predicted transmembrane helix domain (DvhGΔTM) or DvhG with mutated catalytic residue (C317A). b) Quantification of DvhG and the mutated protein levels normalized to total protein loaded. Bars and error bars represent mean and SEM for 3 biological replicates. Representative Western blot images are displayed underneath the bar plot corresponding to the DvhG variants c) Cell surface levels of DvhA-hlx-swap in the presence of 0.2% arabinose. d) DvhA-hlx-swap levels in the culture supernatants at 24 hours in the presence of 0.2% arabinose quantified from dot blots. Statistical analysis was performed using one way ANOVA multiple comparisons (ns, p > 0.05; *, p ≤ 0.05; **, p ≤ 0.01; ***, p ≤ 0.001; ****, p ≤ 0.0001).

To confirm that DvhG was indeed expressed in these strains, we performed Western blots with the strain in the presence of 0.2% arabinose (**Figure 2b**). We observed a band migrating at the expected molecular weight for DvhG (Intensity = 6.85 ± 2.9 X10^3^ a.u.) that was absent in the EV (Intensity = 0.29 ± 0.11 X10^3^ a.u.), indicating that His-DvhG protein is stably expressed under our experimental conditions.

To confirm that these changes in biofilm formation are associated with DvhA-hlx-swap localization, we probed for the adhesin on the cell-surface and supernatants using dot blots. Cell-surface associated dot blots were performed as described in the previous section. To probe for supernatant-localized adhesin, we centrifuged OD_600_ normalized cultures and sterilized the supernatants by passage through a 0.22 µm filter before spotting directly on a nitrocellulose membrane. We observed that the loss of biofilm formation in strains expressing DvhG was associated with a significantly decreased level of DvhA-hlx-swap on the cell-surface (**Figure 2c**) and an increase in the supernatant (**Figure 2d**) compared to the EV control. Taken together, these data suggest that DvhG can process the adhesin leading to a loss of biofilm formed.

As a second confirmation that DvhG-mediated proteolysis results in loss of the DvhA-hlx-swap protein from the cell surface, we targeted the proteolysis site for mutation. Studies from our lab have shown that LapG targets a dialanine site (107TAAG110) on LapA to process this adhesin (30). To test if DvhG targets DvhA-hlx-swap at the di-alanine site 105PAAG108, we mutated both alanine residues to arginine on the chromosome such that the strain will express the mutated adhesin DvhA-hlx-swap (AA-RR). The DvhG expressing plasmid pMQ72-*dvhG* and an empty vector were transformed into the mutated background strain and biofilm formation at 24 hours was compared between both strains, with and without arabinose induction (**Figure S4a**). Without arabinose induction, with only basal levels of DvhG present in the cell, we see robust biofilm formation (OD_550_ = 0.64 ± 0.08) with DvhA-hlx-swap AA-RR mutant. Upon increasing the level of DvhG by arabinose induction, we see a significant decrease in biofilm formation (OD_550_ = 0.40 ± 0.13, p = 0.059; **Figure S4a**), but still above the level of the biofilm formed for a strain carrying the DvhA-hlx-swap with the AA proteolysis site and expressing DvhG (OD_550_ = 0.30 ± 0.03; **Figure 2a**; this level is also represented by the dashed line in **Figure S4a**). In agreement with the biofilm data, in the presence of DvhG, we observed 65% more localization of DvhA-hlx-swap AA-RR on the cell surface and 32% less accumulation in the supernatant than DvhA-hlx-swap (**Figure 2d, S4b, S4c**), suggesting that the mutated adhesin was not processed as efficiently as DvhA-hlx-swap with the WT proteolysis site.

Why did we detect some apparent level of proteolysis in the strain expressing the DvhA-hlx-swap (AA-RR) variant (**Figure S4a**)? It has been previously noted that when LapA was mutated from 108AA109 to 108RR109, LapG was able to access other nearby di-alanine sites on LapA *in vivo* (30). Based on our observations, it is possible that DvhG is also processing other dialanine motifs; dialanines at the N-terminal of LapA and DvhA are highlighted in blue in **Figure S1c**. While the alternative AA target sites of LapA are removed in DvhA-hlx-swap, there are other dialanine motifs further downstream in LapA that could be targeted by DvhG (not shown).

Overall, using multiple lines of evidence, we show that the N-terminus of DvhA has a retention module-like activity that allows localization of the fusion protein to the cell surface, thereby promoting biofilm formation, and that the DvhG-mediated proteolysis site of the DvhA-hlx-swap dialanine motif releases this adhesin from the surface, thereby reducing biofilm formation.

### The role of the predicted transmembrane domain of DvhG

Unlike LapG, which is a periplasmic protease, DvhG has a predicted transmembrane (TM) helix suggesting that this protease may be inner membrane bound, with the catalytic domain localized to the periplasm. To assess whether the TM helix is essential for DvhG proteolytic activity, we cloned *dvhG* lacking the TM helix (deletion of aa 1-40; called *dvhG*ΔTM) with an N-terminal 6x-His tag, into the pMQ72 plasmid (pMQ72-*dvhG*ΔTM). Upon transforming this plasmid into Pf0-1 Δ*lapG*Δ*lapD dvhA-hlx-swap* strain and testing for its ability to form biofilms, we found that this strain was not significantly different from the empty vector control even in the presence of arabinose (OD_550_ = 0.95 ± 0.02 vs 1.02 ± 0.2; **Figure 2a**), indicating that the TM helix is essential to DvhG to process the adhesin. Supporting the biofilm findings, the level of the DvhA-hlx-swap protein on the cell-surface and supernatant for the strain expressing DvhGΔTM was not significantly different from the control (**Figure 2c, 2d**).

The finding above that without its TM helix DvhG cannot cleave DvhA-hlx-swap could be explained by 1) the fact that in the absence of the TM helix the protein is unstable and/or 2) the inability of the variant TM-less DvhG to localize to the periplasm. We do find that the DvhGΔTM protein is more stable in whole-cell lystates than the WT DvhG (Intensity = 149.4 ± 40.0 X10^3^ a.u; **Figure 2b**). Therefore, collectively our findings suggest that the lack of proteolytic activity is likely due to the inability of the variant to access the adhesin in the periplasm, consistent with a lack of a secretion signal in the truncated protein.

### DvhG requires catalytically active cysteine residue to function

Multiple sequence alignment analysis comparing DvhG with LapG-like proteases showed conserved cysteine, histidine, and aspartate (C-H-D) triad that has previously been shown essential for LapG activity (**Figure S3**). To test the importance of the cysteine residue in DvhG, we mutated this residue to alanine (C317A) and cloned the full length mutated *dvhG* into the arabinose-inducible pMQ72 plasmid (pMQ72-*dvhG*-C317A). This plasmid was transformed into Pf0-1 Δ*lapG*Δ*lapD dvhA-hlx-swap* and we compared the biofilm formed, and whole cell and surface levels of the DvhA-hlx-swap protein to the strains containing WT DvhG or the EV.

We performed static biofilm assays with these strains in K10T-1 medium at 24 hours with and without 0.2% arabinose inducer (**Figure 2a**). As expected, the strain expressing DvhG showed reduced biofilm formation when compared to the EV control regardless of the inducer. In contrast, the strain expressing the DvhG-C317A mutant forms a biofilm equivalent to that of the EV control under non-inducing conditions (p = 0.88). With arabinose, DvhG-C317A displays a modest but significant reduction (OD_550_ = 0.69 ± 0.03) in biofilm formation compared to the EV control (OD_550_ = 0.89 ± 0.05), however, these biofilms are still significantly more robust than strain expressing WT DvhG (p < 0.0001). Dot blots also show comparably high levels of DvhA-hlx-swap on the cell surface of EV and DvhG-C317A mutant strain and low levels of the adhesin in their respective supernatants, compared to the strain expressing the WT DvhG (**Figure 2c, 2d**). C317A Western blot shows the protein is as stable of the WT DvhG (Intensity = 6.35 ± 3.8 X10^3^ a.u.; **Figure 2b**).

Our previous studies showed that LapG activity requires calcium ions, consistent with the observation that the structure of LapG contains several calcium ions (29, 35). We also observe calcium-dependence of DvhG (**Figure S4d-f**). Finally, we assessed the ability of LapG to target LapA versus the DvhA-hlx-swap fusion protein, and for DvhG to target these two substrates. These results show that these proteases are proficient in processing their own adhesins but less efficient toward the other adhesins (**Figure S5**).

Together these data suggest that the cysteine residue and calcium ions of DvhG are critical for its proteolytic activity, and that each protease has preference for its cognate proteolysis site.

### DvhD rescues biofilm phenotype in the presence of DvhG

To investigate the effect of the LapD-like DvhD on biofilm formation, we expressed the gene encoding the HA tagged DvhD under an IPTG-inducible P*_tac_* promoter at a neutral *att* site in the Pf0-1 Δ*lapG*Δ*lapD dvhA-hlx-swap* strain along with a *lacI* repressor gene (**Figure 3a**). Fluorescence intensity of a band at the expected ∼75kDa in a whole-cell lysate Western blot showed that DvhD was robustly expressed in the presence of 0.01% IPTG but not without the inducer (**Figure 3b**). We introduced the arabinose-inducible plasmid carrying full length *dvhG* in the Δ*lapG*Δ*lapD dvhA-hlx-swap*/pMQ72-*dvhG* strain background or the Δ*lapG*Δ*lapD dvhA-hlx-swap*/pEV control strain, thus allowing us to control expression of the DvhG and DvhD proteins: 0.2% arabinose to induce DvhG production and 0.01% IPTG to induce DvhD production.

**Figure 3:**
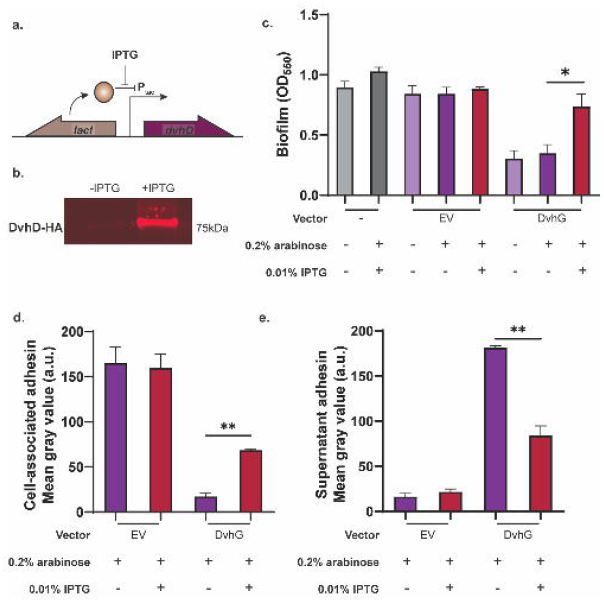
DvhD rescues the biofilm phenotype of Pf0-1. a) The gene encoding HA-tagged DvhD gene from *D. vulgaris* Hildenborough was inserted at the *Tn7 att* site under the control of IPTG-inducible P*_tac_* promoter in the Pf0-1 Δ*lapG*Δ*lapDdvhA-hlx-swap* strain. The *lacI* gene was also introduced at the *att* site to repress *dvhD* gene expression in the absence of IPTG. b) Whole-cell western blot for HA-tagged proteins shows minimal DvhD without IPTG and increased DvhD level in the presence of IPTG. c) Biofilm formed by the Pf0-1 attTn7::*lacI*P*_tac_dvhD*-HA Δ*lapG*Δ*lapDdvhA-hlx-swap* strain in K10T medium at 24 hours. Quantification was performed on the strain containing no vector, empty vector and plasmid expressing full length DvhG. Arabinose at 0.2% and IPTG at 0.01% were used to induce *dvhG* expression from the plasmid and *dvhD* expression from the genome, respectively. d,e) Quantification of DvhA-hlx-swap on the cell surface (d) and the culture supernatant (e) using dot blots in the presence or absence of IPTG. The analyzed strains carried either an empty vector or a plasmid expressing full length DvhG, all under inducing conditions with 0.2% arabinose. Statistical analysis was performed using one way ANOVA multiple comparisons (ns, p > 0.05; *, p ≤ 0.05; **, p ≤ 0.01; ***, p ≤ 0.001; ****, p ≤ 0.0001).

We performed static biofilm assays under three conditions – without any inducer, with arabinose only (expressing DvhG only), and with both arabinose and IPTG (expressing both DvhG and DvhD; **Figure 3c**). We compared these data with an empty vector control under the same conditions. In the arabinose only control, we observed a decrease in biofilm consistent with DvhG-dependent proteolytic activity. However, in the presence of arabinose and IPTG, when DvhD is expressed, we noted up to 56% increase in biofilm compared to the arabinose-only control. Under these latter conditions the biofilm formed was on par with the no vector and EV positive controls. The dot blot analysis shows that in the presence of both arabinose and IPTG, DvhA-hlx-swap recovered on the cell-surface by 75% compared to the arabinose-only condition, and consequently, the supernatant associated DvhA-hlx-swap decreased by 54% compared to arabinose-only condition (**Figure 3d, e**). These data suggest that the presence of DvhD rescues biofilm phenotype, likely by impacting the ability of DvhG to target its substrate.

### Biofilm recovery mediated by DvhD is driven by c-di-GMP levels

For Pf0-1, a low concentration of extracellular inorganic phosphate (Pi) has been shown to reduce LapA localization to the cell surface (36). Low Pi activates the production of c-di-GMP PDE RapA, which decreases intracellular c-di-GMP levels and in turn reduces the pool of c-di-GMP to bind to LapD, and eventually, loss of LapA from the cell surface. To test if the rescue in biofilm formation by expressing DvhD is driven by cellular c-di-GMP levels, we manipulated intracellular c-di-GMP concentrations by modulating Pi concentration in K10T-1 medium.

We performed static biofilm assays with both DvhD and DvhG induction, DvhG-only induction and EV control in K10Tπ medium (low phosphate medium, ∼0.1 μM; associated with low c-di-GMP) or with high extracellular phosphate (K10T medium, 1 μM, associated with relatively high c-di-GMP). Using these experimental conditions, we calculated the “Δ” which represents the difference in biofilm formed by the strain expressing both DvhG and DvhD (induction with both arabinose and IPTG) and biofilm formation by the strain expressing DvhG but not DvhD (induction with arabinose only). Our results indicate that Δ for high Pi condition is 0.5 ± 0.14, which is significantly larger than low Pi condition 0.12 ± 0.044 (**Figure 4a, S6a**). These data indicate that shifting the Pi concentration (and thus c-di-GMP levels) can impact DvhD/G-mediated biofilm formation.

**Figure 4:**
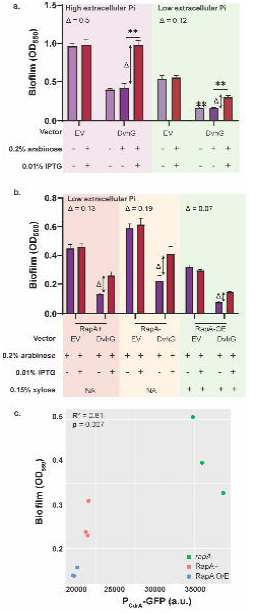
Biofilm rescue by DvhD is c-di-GMP dependent. a) Biofilm formed in the Pf0-1 attTn7::*lacI*P*_tac_dvhD*-HA Δ*lapG*Δ*lapDdvhA-hlx-swap* strain in low phosphate K10Tπ medium (∼0.14 mM) or high phosphate K10T medium (1 mM) at 24 hours. The strain contains either no vector, an empty vector or a plasmid expressing full length DvhG. Arabinose at 0.2% concentration and IPTG at 0.01% concentration are used to induce *dvhG* expression from the plasmid and *dvhD* expression from the genome, respectively. “Δ” quantifies the extent of biofilm rescue by DvhD and is represented by the difference in biofilm formation by the strain expressing DvhG but not DvhD (purple bar) and biofilm formation by the strain expressing both DvhG and DvhD (red bar). b) Biofilm formed by the Pf0-1 attTn7::*lacI*P*_tac_dvhD*-HAΔ*lapG*Δ*lapDdvhA-hlx-swap* strain in low phosphate K10Tπ medium at 24 hours with, active phosphodiesterase RapA (RapA^+^), without RapA (Δ*rapA*) or with RapA overexpressed by xylose-inducible P_xutR_ promoter (RapA OE) in the background. The extent of biofilm rescue by DvhD “Δ” is compared among the three strains and indicated over the bar plots. Statistical analysis was performed using one way ANOVA test (ns, p > 0.05; *, p ≤ 0.05; **, p ≤ 0.01; ***, p ≤ 0.001; ****, p ≤ 0.0001). c) Graph representing intracellular c-di-GMP levels calculated from GFP fluorescence from a P_CdrA-_*gfp* promoter transcriptional fusion plotted versus biofilm formed when DvhD is induced. Linear correlation (R^2^) and statistical significance was calculated using Pearson’s test in R (v 4.3.0).

Next, we made a markerless *rapA* deletion mutation yielding the Pf0-1 att::*lac*IP*_tac_dvhD*-HA Δ*lapG*Δ*lapD*Δ*rapA dvhA-hlx-swap* strain. In low Pi conditions, when RapA is activated and c-di-GMP is hydrolyzed, a Δ*rapA* strain should not affect the c-di-GMP levels. Conversely, we also constructed a strain in which we placed *rapA* under the control of a xylose-inducible promoter. Induction with xylose under low Pi conditions is expected to increase the level of RapA and drive the intracellular c-di-GMP levels lower than WT. We transformed pMQ72-*dvhG* and EV in these backgrounds and tested for biofilm formation in the low Pi medium and compared conditions with both DvhD and DvhG induction, DvhG-only induction, and EV control (**Figure 4b**). RapA overexpression was induced with 0.15% xylose.

While we consistently observed higher Δ values in the biofilm formed by the *rapA* deletion strain (0.19 ± 0.11) than in the strain with an active RapA (0.13 ± 0.02), these differences were not significantly different (**Figure 4b, S6b**). On the other hand, we observed lower Δ values when RapA was produced under the control of the xylose promoter (0.07 ± 0.01) compared to when WT RapA levels were produced. However, the difference was also not significant.

We then quantified the levels of intracellular c-di-GMP using a P*_cdrA_*promoter fused to GFP (39) (**Figure 4c**). We observe a significant positive association between c-di-GMP level and biofilm formed (R^2^= 0.81; p = 0.007). These data show that altering c-di-GMP in this heterologous background results in the expected outcome (low c-di-GMP➔low biofilm, hi c-di-GMP➔high biofilm) for a c-di-GMP-responsive system.

## Discussion

In this study, we characterize the posttranslational regulation of a large adhesive protein important for biofilm formation in *Desulfovibrio vulgaris* Hildenborough (DvH). Our bioinformatic analysis along with work from De Leon et al. revealed T1SS machinery, LapD- and LapG-like proteins and large adhesins, similar to the Lap system of *P. fluorescens* Pf0-1 that is critical for biofilm formation.

Here we find that certain aspects of the two Lap systems are functionally similar. The large adhesin DvhA appears to have a retention module at its N-terminus and is predicted to be comprised of folded beta sheets. The adhesin also shares a highly conserved dialanine site with LapA, which acts as a target site for proteolysis by DvhG. Like LapG, DvhG is a calcium dependent cysteine protease that cleaves the large adhesin, leading to a loss of biofilm. Further, we identified a protein DvhD, that rescues biofilm formation even in the presence of DvhG. In the Pf0-1 Lap system, LapD, which seems to be functionally analogous to DvhD, also rescues biofilm formation (24). Further structural and biochemical studies have shown that LapD can sequester LapG via a conserved tryptophan (Trp) residue, thus preventing the proteolysis of LapA, and rescuing the biofilm (40, 41). While we do not have direct evidence yet showing that DvhG interacts with DvhD, we hypothesize that this interaction could be responsible for the biofilm rescue phenotype despite the very different periplasmic domains of LapD and DvhD, the latter of which lacks a conserved Trp residue. Lastly, we also determined that intracellular levels of c-di-GMP contribute to regulating the Lap machinery of DvH, although as outlined below, with some likely differences from the Pf0-1 system.

While there are broad functional similarities between the Pf0-1 and DvH Lap system, we also note some interesting differences between the two systems. Our results indicate that the proteases LapG and DvhG are highly specific towards their native adhesins and show limited cross-reactivity. This finding suggests that the way in which LapA-LapG interact could be distinct from DvhA-DvhG interaction. Sequence analysis of the N-terminus of LapA-like adhesins show a conserved residue pair “DP” in addition to the dialanine motif (22), however, these residues are absent in DvhA. While the function of these residues has not been explicitly tested, we hypothesize that they may be responsible for imparting the adhesins structural specificity towards the proteases.

We hypothesize that the interaction between DvhG and DvhD is unique. As mentioned earlier, in Pf0-1 LapG interacts with LapD via a highly conserved tryptophan residue in the periplasmic domain of LapD (40, 41). This residue inserts itself into a hydrophobic binding pocket formed on the surface of LapG. However, sequence analysis of DvhD shows no conserved tryptophan residue. This suggests the presence of new distinct interface/residues involved in the DvhG and DvhD interaction. We also note that unlike the periplasmic LapG, DvhG is likely an inner membrane bound protein, which could also affect the mechanics of this interaction.

Furthermore, unlike LapD which has an inactive, cytoplasmic GGDEF-EAL domain, DvhD consists of a cytoplasmic, inactive HD-GYP domain. As DvH is an environmental microbe belonging to the class *deltaproteobacteria,* which are known to have higher fraction of HD-GYP domain containing proteins (42), perhaps this SRB has re-purposed one if this family member as a c-di-GMP receptor.

Finally, the c-di-GMP network of DvH appears to be complex with 40 DGCs, PDEs and dual domain proteins predicted through sequence analysis (43). This is almost on par with the Pf0-1 c-di-GMP circuit with 51 proteins (44). However, given the difference in numbers and domains of c-di-GMP sensing proteins (different substrate-binding mechanics/affinities/dynamics) in the two microbes, the dynamics of c-di-GMP regulation, and therefore, the intracellular concentrations of this second messenger could be different. Consistent with this idea, our experiments indicate that DvhD is only partially responsive to changing c-di-GMP levels in Pf0-1. It is worth noting that most of our experiments performed here manipulating c-di-GMP levels were done in low phosphate conditions. While we know that c-di-GMP levels increase in a Δ*rapA* strain under low phosphate, the lack of phosphate is likely affecting other physiological processes that could hamper biofilm formation (36). Furthermore, data from Font, Sondermann et al. (personal communication) show that the affinity of the cytoplasmic PAS-HDGYP domain of DvhD for c-di-GMP in the high nanomolar range. Previously measurements with LapD show affinity to be in the low micromolar range (24, 25). Thus, one caveat of our study is that heterologous expression of DvhD in Pf0-1 may be in the context of a higher range of c-di-GMP concentrations, making it more challenging to effectively vary concentrations in the physiological range of DvhD.

Our future work will explore the differences between the two Lap systems. We are especially interested in finding the residues that facilitate binding between DvhG and DvhD, and DvhA and DvhG. Furthermore, we aim to explore whether the c-di-GMP interaction network among the multiple DGCs and LapD that controls biofilm formation by Pf0-1 also exists in DvH. Finally, beyond DvH, we plan to use Pf0-1 as a chassis to study Lap-like systems of other microbes that may be hard to culture and/or be genetically intractable to broaden our understanding of biofilm formation via the Lap system.

## Materials and Methods

### Strains and growth conditions

*Pseudomonas fluorescens* Pf0-1 Δ*lapG*Δ*lapD* strain (23) was used as a base strain in the studies here. Mutations were made in this background using *E. coli* S17-1 λpir. Strains used in this study are listed in Supplementary Table 1. Bacteria were cultured in lysogeny broth (LB) or plated on 1.5% LB agar with antibiotics as needed at 37°C for *E. coli* and at 30°C for *P. fluorescens* Pf0-1 (abbreviated Pf0-1). Gentamycin (Gm) was used at a concentration of 10 µg/ml for *E. coli* and at 30 µg/ml for Pf0-1. Carbenicillin (Cb) was used at 100 µg/ml for *E. coli*. Tetracycline (Tet) was used at 15 µg/ml for *E. coli* and 30 µg/ml for Pf0-1. All assays were performed in K10T-1 medium (45): Tris buffer, pH 7.4 (50 mM), Bacto tryptone (0.2% (w/v)), glycerol (0.15% (v/v)), K_2_HPO_4_ (1 mM) and MgSO_4_ (0.61 mM). For phosphate limited conditions, K_2_HPO_4_ was excluded from K10T-1 medium, yielding K10Tπ. The medium was assembled from concentrated stock solutions that were either autoclaved (121°C, 15 minutes) or filter sterilized. For calcium chelation, K10T-1 was supplemented with 40 µM ethylene glycol-bis(2-aminoethylether)-N,N,N’,N’-tetraacetic acid (EGTA). Genes under the control of the P_BAD_ promoter were induced with arabinose (0.2% (w/v)) and those under P_tac_ promoter were induced with isopropyl-D-thiogalactopyranoside (IPTG, 0.5 mM).

### Construction of plasmids and mutant strains

All plasmids and primers used in this study are listed in Supplementary Table 1 and Table 2, respectively. Inserts for plasmid construction were amplified from *Desulfovibrio vulgaris* (strain ATCC 29579 / DSM 644 / NCIMB 8303 / VKM B-1760 / Hildenborough) genome. All vectors were assembled with the inserts using Gibson assembly (46). The *lapA*-*dvhA* adhesin fusions (*dvhA-RM-swap* and *dvhA-hlx-swap*) and *dvhA-hlx-swap* 105PRRG108 were built by allelic exchange using pMQ30 plasmid as described (46). Expression vectors were built using arabinose-inducible pMQ72 vector with genes placed under P_BAD_ promoter. Insertion at the neutral *att* site of Pf0-1 were built using pMQ56 mini-*Tn7* vector with the helper plasmid pUX-BF13. To remove the resistance marker, pFLP3 plasmid was used, followed by sucrose counterselection. Mutations in *dvhD* gene at Pf0-1 *att* site were introduced by allelic exchange using pMQ30 plasmid. Pf0-1 *lapB::*pMQ89 was constructed by single-crossover, conferring gentamycin resistance to the strain. All plasmids were constructed in *E. coli* and introduced into Pf0-1 by electroporation, as reported (38).

### Static biofilm assays

Static biofilm assays were performed as previously described in (36) in K10T-1 or K10Tπ media at 24 hours at 30°C using crystal violet staining. Details of the procedure are described in SI text.

### Quantitative adhesin surface localization assay using dot blots

Localization of the adhesin was probed with the help of dot blots using Pf0-1 strains with a functional 3xHA-tagged adhesin as described in (36) Briefly, overnight Pf0-1 cultures were subcultured in 5 ml K10T-1 medium (1:100 dilution) and incubated for 24 hours. The cultures were OD_600_ normalized and spotted directly on nitrocellulose membrane. Supernatant samples were prepared by filter-sterilizing the OD_600_ normalized cultures before spotting on the membrane. The blots were probed for the adhesin using chemiluminescence.

### Detection and quantification of proteins using Western blots

Pf0-1 strains with genome-integrated DvhD harboring a functional HA tag or with adhesins harboring 3x-HA tag were used to detect and quantify the proteins. The samples were prepared and processed as described above for dot blots. Pf0-1 strains with 6x-His tagged DvhG (or DvhG variants) on a pMQ72 plasmid were used to detect and quantify this protein. Empty vector pMQ72 was used as a negative control. For all Western blots, subcultured samples were normalized for cell density (OD_600_) and protein content using BCA assay (Pierce). After resolving the samples on SDS-Page gels and transferring to a nitrocellulose membrane, the proteins were quantified using fluorescence intensity. Detailed procedures of Western blots are provided in the SI text.

### Bioinformatics analysis

The Simple Modular Architecture Research Tool (SMART, http://smart.embl-heidelberg.de/) was used to identify LapG- and LapD-like proteins. LapG contains a “Pfam: Peptidase_C93” domain and LapD-like proteins contain a perpplasmic “Pfam: LapD_MoxY_N” domain. The SMART Domain selection tool was used to identify proteins in the database containing either domain and the FASTA results were saved for analysis in R. Bacterial species encoding LapG-like and LapD-like were identified using the R “intersect” function and genomes encoding only LapG or LapD were identified using the “setdiff” function. The Bacteria and Viral Bioinformatic Resource Center (https://www.bv-brc.org/) was used to manually inspect the genomic regions surrounding LapG.

### Statistics

Data visualization and statistical analysis were performed in GraphPad Prism 9 (v.9.2.0). Linear models were built in R (v.4.3.0) and visualized using ggplot2 (v.3.4.2). The script used to perform the analyses can be found at https://github.com/GeiselBiofilm.

## Acknowledgements.

We would like to thank Prof. Will D. Leavitt for providing us with the *Desulfovibrio vulgaris* strain used in this study and helping us kick-start the project. We thank Dr. Thomas Hampton and Kaitlyn Barrack for their help with statistical analysis. We also thank Dr. Sherry Kuchma and Christopher Geiger for thoughtful discussions and help with flow cytometry. This work was supported by NIH grant R01GM123609 G.A.O. This work was also supported by bioMT through NIH NIGMS grant P20-GM113132.

