## Supplemental Material for "Reconstitution of a Biofilm Adhesin System from a Sulfate-Reducing Bacterium in *Pseudomonas fluorescens*"

### Supporting Information

#### Figures

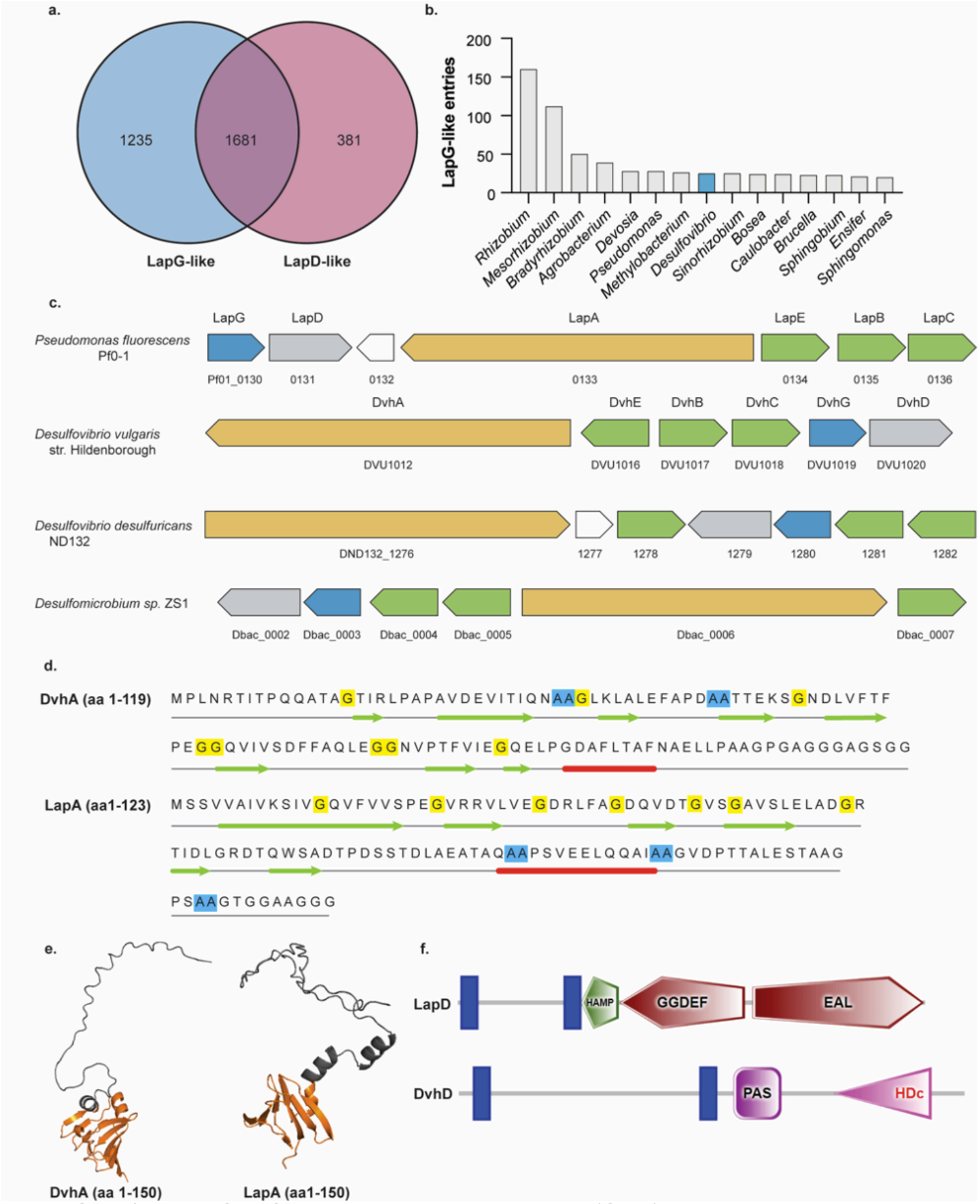


**Figure S1**: a) The NCBI Conserved Domain Database (CDD) was used to gather organisms encoding LapG-like (pfam06035) and LapD-like (pfam16448) proteins. Each group was separated into a list and the R programming language package VennDiagram was used to find the overlap between the groups. b) Counts of LapG-like proteins in various microbial genera. Proteins belonging to the genus *Desulfovibrio* are highlighted in blue, ABC transporters in green, and adhesins in yellow. c) Genomic arrangement of Lap components of *P. fluorescens* Pf0-1, *D. vulgaris* Hildenborough, *D. sulfuricans* ND132 and *Desulfomicrobium sp.* ZS1. LapG-like proteins are highlighted in blue. d) Secondary structure prediction using JPred4 of DvhA aa 1-121 and LapA aa1 -123. Green arrows represent beta sheets, red arrows represent alpha-helix. Double alanine motifs are highlighted in blue. e) AlphaFold structure prediction of the first 150 amino acids of LapA and DvhA displaying a folded globular domain at the N-terminus of the two proteins (shown in orange), likely serving as a retention module. f) Comparison of domain architectures of LapD (top) and DvhD (bottom) and using Simple Modular Architecture Research Tool (SMART).

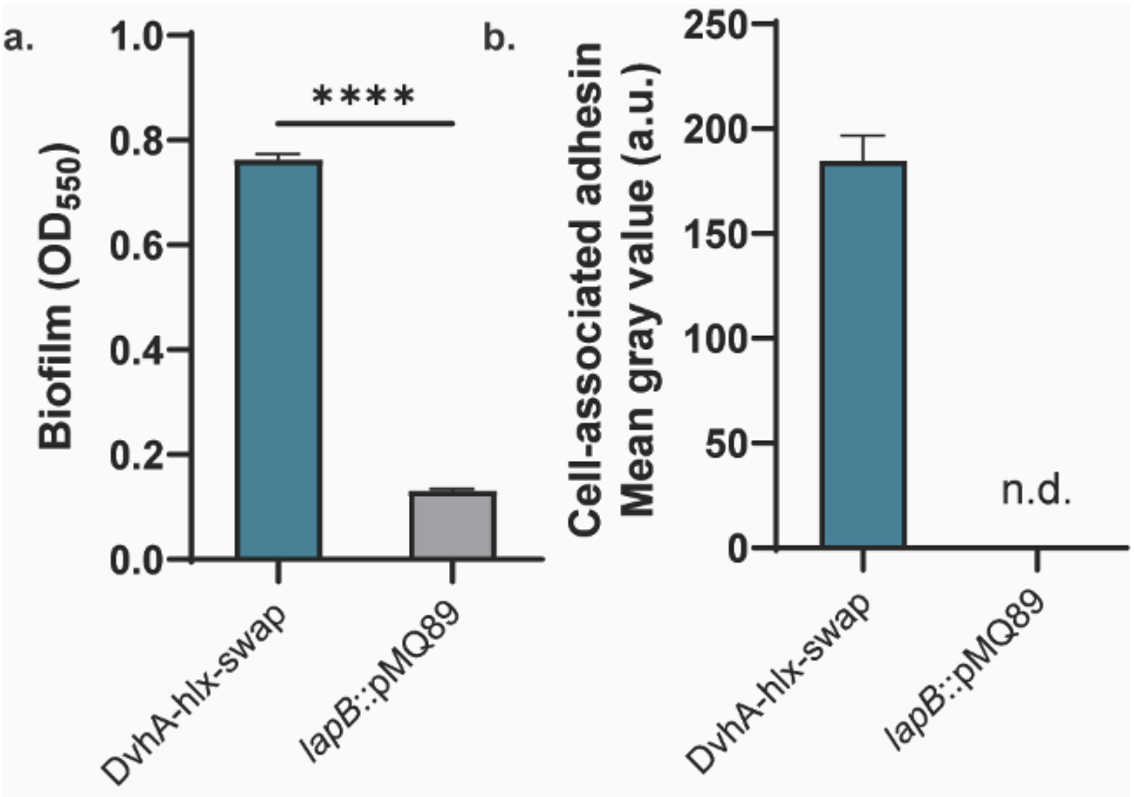


**Figure S2**: DvhA-hlx-swap used T1SS machinery of *P. fluorescens* Pf0-1. a) Biofilm formation in K10T-1 medium at 24 hours with DvhA-hlx-swap with and without the Pf0-1 T1SS ABC transporter component LapB. b) Quantification of cell-surface associated adhesin in both strains using dot blots. Signal intensity for lapB::*pMQ89* was not detected (n.d.) Statistical analysis was performed using unpaired two-tailed t-test (ns, p > 0.05; *, p ≤ 0.05; **, p ≤ 0.01; ***, p ≤ 0.001; ****, p ≤ 0.0001).

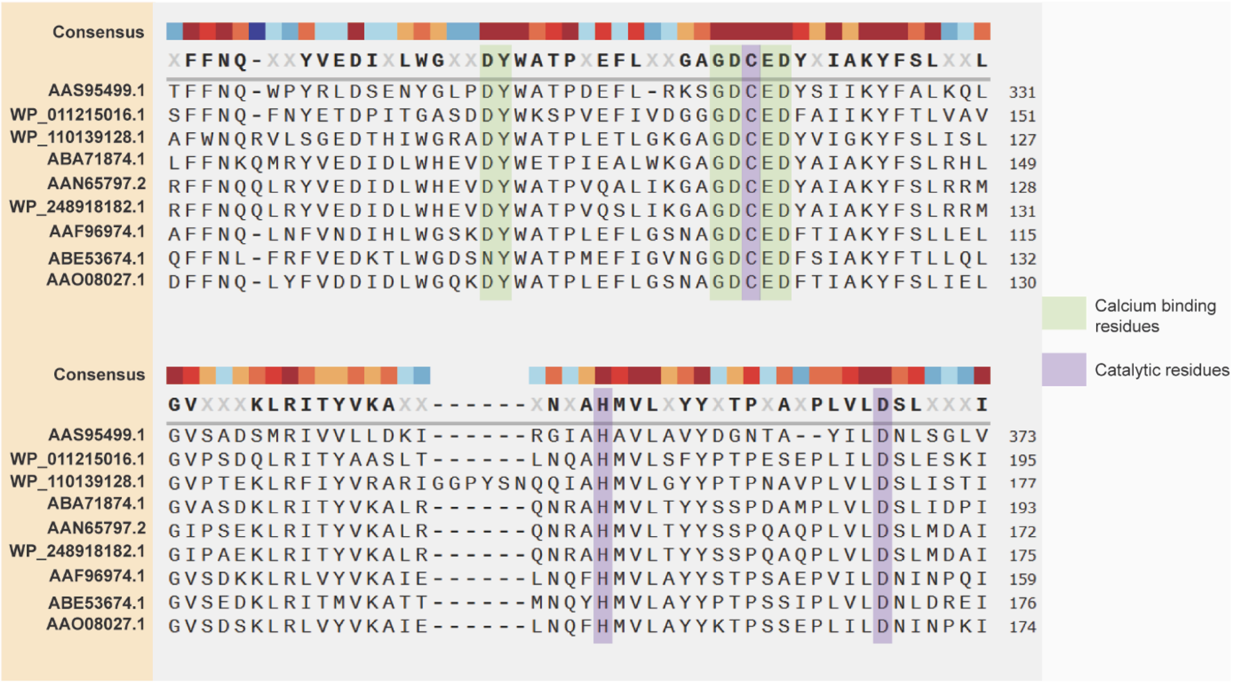


**Figure S3**: Multiple Sequence Alignment of DvhG with LapG-like proteases. Multiple sequence alignment of DvhG with LapG-like proteases using MUSCLE reveals conserved catalytic triad C-H-D (purple) and conserved calcium binding residues (green). The sequences used in the alignment are LapG-like proteases from *Desulfovibrio vulgaris* Hildenborough, *Legionella pnemophila, Bordetella bronchiseptica, Pseudomonas fluorescens* Pf0-1, *Pseudomonas putida* KT2440, *Pseudomonas entomophila, Vibrio cholerae* O1 biovar El Tor str N16961, *Shewanella denitrificans* OS217 and *Vibrio vulnificus* CMCP6, respectively.

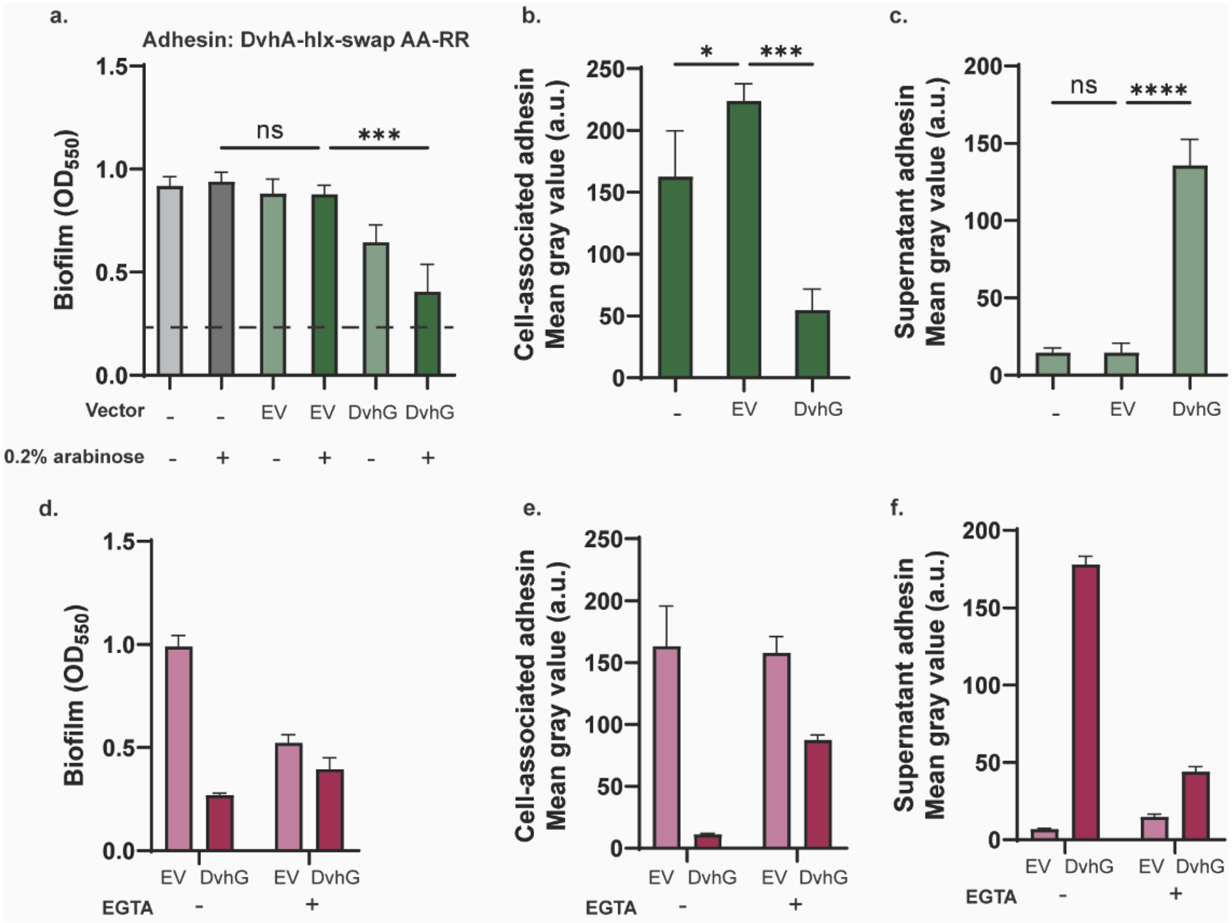


**Figure S4**: DvhG is a calcium ion dependent protease that targets dialanine in DvhA-hlx-swap. a) Biofilm formed in K10T-1 medium at 24 hours with *dvhA-hlx-swap* mutated at the dialanine site from 105PAAG108 to 105PRRG108 without a vector, with empty vector (EV) and *dvhG* expressed on an arabinose inducible plasmid in the presence or absence of arabinose. The dashed line represents the mean biofilm formed when DvhG processes the unmutated dialanine. b) Cell surface levels of DvhA-hlx-swap AA-RR in the presence of 0.2% arabinose. c) DvhA-hlx-swap AA-RR levels in the culture supernatants in the presence of 0.2% arabinose. d) Biofilm formed in K10T medium at 24 hours by the Pf0-1 Δ*lapG*Δ*lapDdvhA-hlx-swap* strain with arabinose inducible empty vector (EV) or vector expressing dvhG in the presence of 0.2% arabinose, with or without 40 μM EGTA (a calcium chelator). e) Quantification of cell-surface associated DvhA-hlx-swap with EV or *dvhG* induced with arabinose, with or without EGTA. f) DvhA-hlx-swap levels associated with the culture supernatants of the corresponding strains. Statistical analysis was performed using one way ANOVA multiple comparisons (ns, $p > 0.05$; *, $p \leq 0.01$; ***).

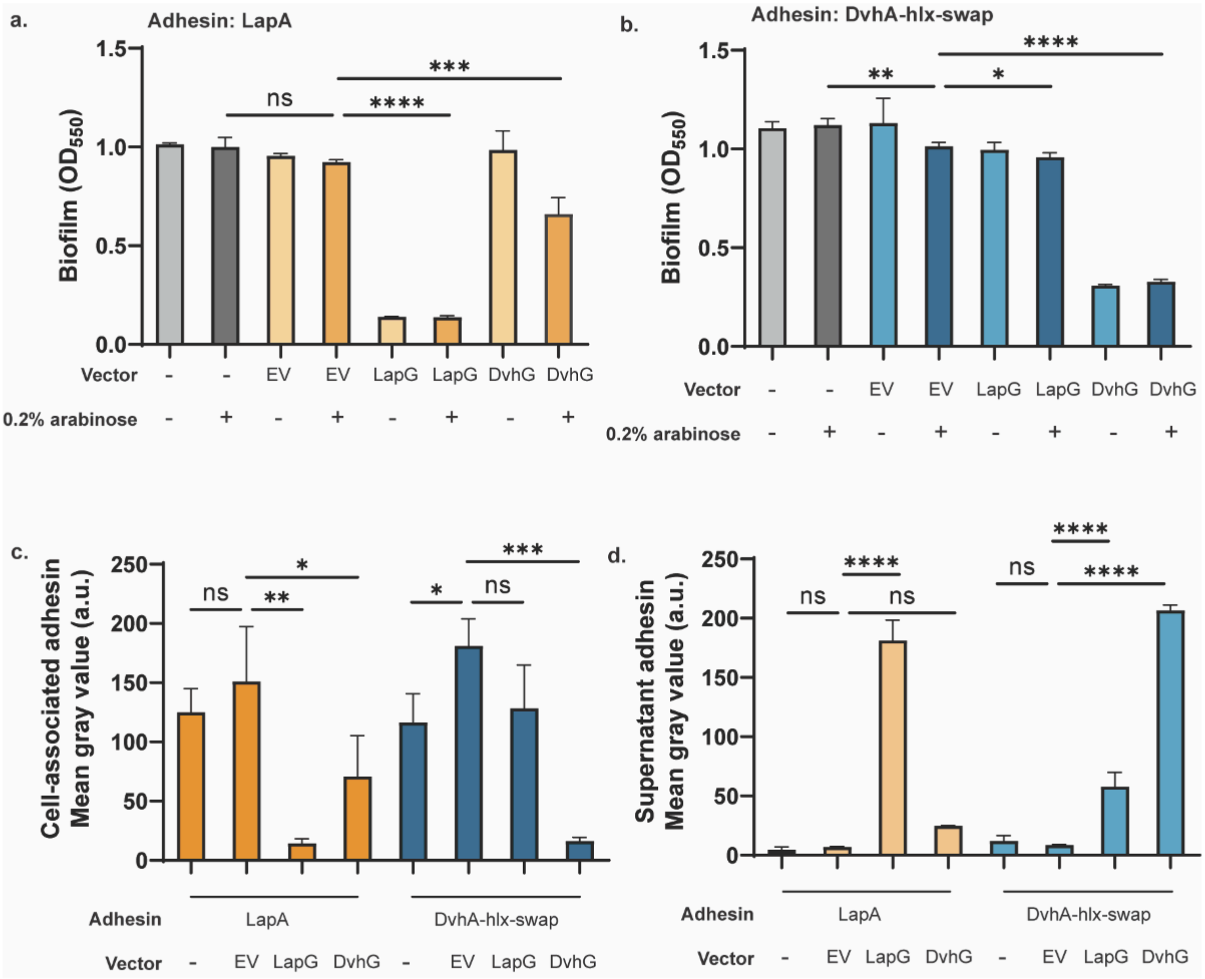


**Figure S5**. LapG and DvhG display specificity towards their native adhesins. a) Biofilm formation in K10T medium at 24 hours by the Pf0-1 Δ*lapG*Δ*lapD* strain expressing the native adhesin LapA. Quantification was performed on the strains carrying no vector, arabinose inducible empty vector (EV), plasmid expressing *lapG* or *dvhG* in the presence or absence of arabinose. b) Biofilm formation in K10T medium at 24 hours by the Pf0-1 Δ*lapG*Δ*lapD* strain expressing the fusion adhesin DvhA-hlx-swap. Quantification was performed on the strains carrying no vector, arabinose inducible empty vector (EV), plasmid expressing *lapG* or *dvhG* in the presence or absence of arabinose. c) Quantification of cell-surface associated adhesin of the strains in the presence of 0.2% arabinose. The bars in orange represent LapA and the bars in teal represent DvhA-hlx swap. d) Quantification of adhesin levels associated with the culture supernatants of the strains in the presence of 0.2% arabinose. The bars in coral represent LapA and the bars in light blue represent DvhA-hlx swap. Statistical analysis was performed using one way ANOVA multiple comparisons against the empty vector control (ns, $p > 0.05$; *, $p \le 0.05$; **, $p \le 0.01$; ***, $p \le 0.001$; ****, $p \le 0.0001$).

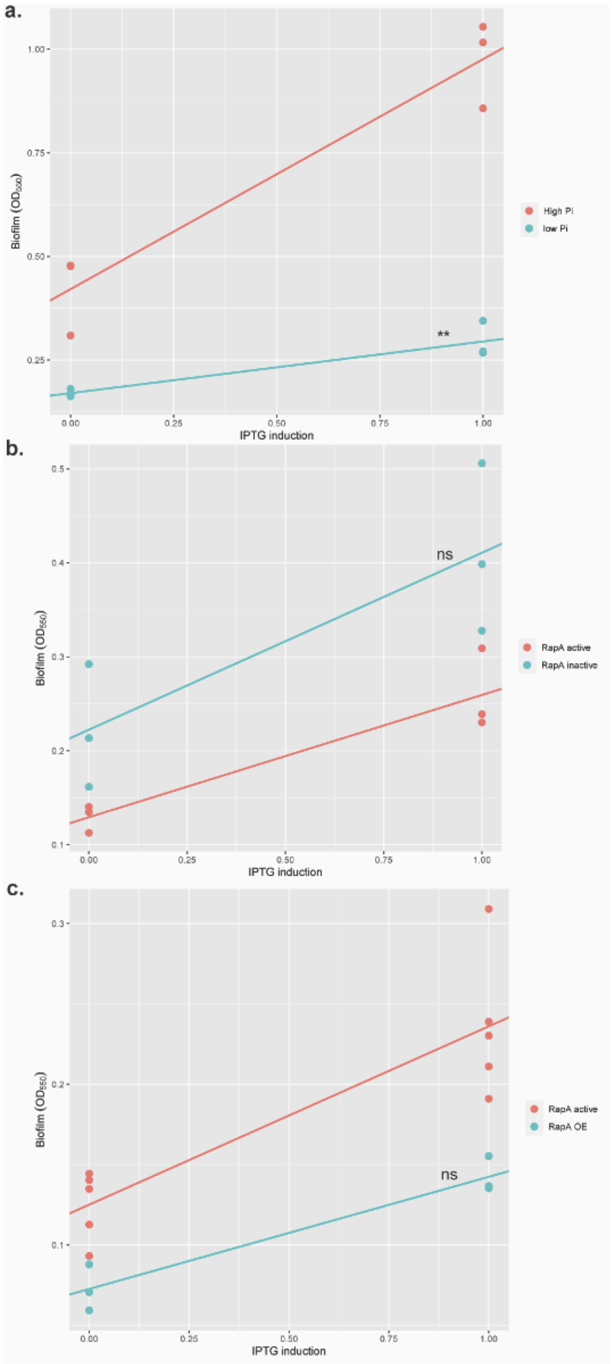


**Figure S6**: Statistical analysis to determine if biofilm rescue is dependent on c-di-GMP. Plots showing the effect of IPTG induction (0 – no IPTG, 1- 0.01% IPTG) on biofilm formation under a) high phosphate (red) vs low phosphate (blue) conditions. b) RapA-active (red) vs RapA-inactive (blue) strain under low phosphate conditions c) RapA-active (red) vs RapA overexpression (RapA OE, blue) strain under low phosphate conditions. Significance in all cases was calculated by linear regression with interaction term in R language (https://github.com/GeiselBiofilm/).

**<u>SI Text</u>**

**Proteolytic activity of DvhG depends upon calcium ions.**

Calcium has been shown to influence biofilm formation by bacteria (1, 2). For *P. fluorescens* Pf0-1, low calcium concentrations result in enhanced biofilm formation, as this lack of calcium negatively affects LapG's Ca-dependent activity and thus proteolysis of the N-terminus of LapA (3). The Multiple Sequence Alignment analysis also revealed conserved calcium-binding sites that could be critical for proteolysis (3, 4) (**Figure S3**).

To test if the activity of DvhG is dependent upon calcium ions, we added a calcium chelator (EGTA) to the K10T-1 medium to sequester extracellular calcium (5). Static biofilm assays were performed at 24 hours. We compared the biofilm formed by strain Pf0-1 *ΔlapGΔlapD dvhA-hlx-swap* expressing DvhG to the strain harboring the empty vector under conditions with and without 40 µM EGTA, all in the presence of arabinose (**Figure S4d**). We found that EGTA compromised the ability of even our EV control to form biofilms. Upon calculating the percentage decrease in biofilm formation in the presence or absence of EGTA between the EV and DvhG, we find that this change is much smaller in the presence of EGTA (29%) than the absence (68%).

The dot blot analysis showed us a more convincing picture. Despite the observed reduction in biofilm formation +EGTA, EV controls under EGTA- and EGTA+ conditions were not significantly different ($p = 0.88$; **Figure S4e**), suggesting that the defect in biofilm formation in EV controls was due to a factor besides DvhG activity. For example, RTX proteins like LapA require Ca-binding to facilitate folding (6). Finally, we noted that 92% reduction in adhesin level on the cell surface when expressing DvhG without EGTA, but only 44% reduction with EGTA, implying the added EGTA reduced DvhG activity. The supernatant dot blots also complement these data (**Figure S4f**). Together, these data are consistent with the conclusion that calcium is required for the DvhG proteolytic activity.

**LapG and DvhG display specificity towards their native adhesins.**

To test whether LapG and DvhG can proteolyze their non-native adhesins, static biofilm assays were conducted at 24 hours comparing a plasmid carrying full length *lapG* (pMQ72-*lapG*), a plasmid carrying full length *dvhG* (pMQ72-*dvhG*) and an empty vector in two background strains: Pf0-1 *ΔlapGΔlapD* which expresses WT LapA adhesin and *ΔlapGΔlapD dvhA-hlx-swap* which expresses the fusion protein DvhA-hlx-swap. The plasmids were induced with 0.2% arabinose and compared to un-induced conditions.

Our results show that, when LapG is present, the strain expressing WT LapA has a significantly reduced ability to form a biofilm. With DvhG, on the other hand, the biofilm phenotype is significantly reduced as compared to the EV in the presence of arabinose ($OD_{550}$ = 0.66 ± 0.08 vs 0.98 ± 0.09, p = 0.01), however it is also significantly higher than biofilms formed with a strain expressing LapG ($OD_{550}$ = 0.13 ± 0.008, p = 0.0004; **Figure S5a**).

When LapG is induced with arabinose in the strain expressing DvhA-hlx-swap, we observe a significant but modest loss in biofilm formation compared to the EV control ($OD_{550}$ = 0.95 ± 0.02 vs 1.01 ± 0.02). With DvhG expressed in this background, as expected, the loss of biofilm phenotype was more striking in comparison to the EV control (**Figure S5b**).

We calculated the percentage decrease in biofilm formation from the EV control with both proteases in arabinose-induced conditions. We noted that LapG caused an 85% decrease with LapA but only 6% decrease with DvhA-hlx-swap. DvhG caused a 68% decrease with DvhA-hlx-swap but only 30% decrease with LapA. Dot blot analysis suggests that changes in biofilm phenotype are due to changes adhesin levels on the cell surface, with a concomitant increase in the levels of adhesin in the supernatant (**Figure S5c-d**). These results suggest that these proteases are proficient in processing their own adhesins but less efficient towards the other adhesin.

#### Additional Materials and Methods

##### Static biofilm assays.

Pf0-1 strains were struck on LB agar plates overnight and a single colony was used to inoculate 5 ml LB for 16-18 hours of growth. 1.5 µl of the LB overnight culture was used to inoculate 100 µl of K10T-1 (or K10Tπ) medium in a 96-well round bottom polypropylene plates (Corning Life Sciences, Glendale, AZ). The plate was incubated in a humidified box at 30ºC for 24 hours. The planktonic cells were discarded, and the plate was washed with water to remove any loosely bound cells. To stain the biofilms, 125 µl crystal violet (0.1% (w/v)) was added to the wells and incubated at room temperature for 20 minutes. Excess stain was washed-off twice with water and the wells were allowed to dry before de-staining with 150 µl solution of water, methanol, and glacial acetic acid (45:45:10) for 5 minutes. 100 µl of the solubilized crystal violet from the wells was transferred to a 96-well flat bottom plate and the optical density was measured at 550 nm wavelength. Solubilized crystal violet from an uninoculated well was used as a blank for the measurements.

**Quantitative surface adhesin localization assays using dot blots.**

Pf0-1 strains expressing a WT 3x HA-tagged LapA of LapA variant were struck on LB agar plates overnight and a single colony was used to inoculate 5 ml LB for 16-18 hours. The LB overnight culture was used to inoculate 5 ml of K10T-1 medium (1:100) for growth at 30ºC for 24 hours. The cultures were normalized to an $OD_{600}$ value of 0.5 and pelleted by centrifugation (13,200 rpm, 1 minute). The supernatants were collected, and filter sterilized using 0.22 µm syringe filters to probe for supernatant associated LapA. The pellets were washed with K10T-1 medium twice by resuspension and centrifugation. The pellets were finally resuspended in K10T-1 medium to an $OD_{600}$ of 0.5 to probe for cell-surface associated LapA. 5 µl of cell-surface associated and supernatant fractions were spotted on 0.2 µm nitrocellulose membrane (Bio-Rad Laboratories, Hercules, CA) and allowed to dry completely. The membranes were then incubated in a blocking solution containing 3% bovine serum albumin (BSA) in Tris-buffered saline with 1% Tween 20 (TBST) for 1 hour at room temperature. The blots were probed for LapA-HA using anti-HA antibody (BioLegend, San Diego, CA) at 1:2000 dilution in TBST containing 3% BSA overnight at 4ºC. The membranes were washed thrice in TBST to remove excess antibody and incubated in horseradish peroxidase conjugated anti-mouse secondary antibody (1:15,000 dilution) in TBST for 1 hour at room temperature. Next, the membranes were washed twice in TBST and once in Tris-buffered saline (TBS) and incubated with Western Lightning Plus-ECL enhanced chemiluminescence substrate (PerkinElmer, Waltham, MA) for 1 minute. The membranes were imaged on a BioRad ChemiDoc MP Imaging System. Images were quantified using ImageJ software (NIH) as previously described in (7)

**Detection and quantification of the proteins using Western blots.**

Pf0-1 strains of interest were struck on LB agar plates (with antibiotics, if needed) overnight and a single colony was used to inoculate 5 ml LB for 16-18 hours of growth. The LB overnight culture was used to inoculate 5 ml of K10T-1 medium (1:100) at 30ºC for 24 hours. DvhG overnights were subcultured (1:100) in 50 ml K10T-1 media. The cultures were $OD_{600}$ normalized and pelleted by centrifugation (13,200 rpm, 1 minute). The pellets were resuspended, lysed using French press and quantified for protein using Pierce™ BCA assay (ThermoFischer Scientific, Waltham, MA). The samples were boiled for 5 minutes in Laemmeli sample buffer with 2-mercaptoethanol and were resolved on 7.5% (for adhesins) or 10% (for DvhD and DvhG) polyacrylamide gels. After transferring the proteins to a 0.2 µm nitrocellulose membrane (Bio-Rad Laboratories, Hercules, CA) and blocking with 3% BSA in TBST for 1 hour, the proteins were incubated in mouse anti-HA antibody in TBST (1:2000 dilution) overnight at 4ºC for DvhD and adhesins. For DvhG, the blots were incubated in Invitrogen mouse anti-His antibody at the same dilution (ThermoFischer Scientific, Waltham, MA). Fluorescently labelled anti-mouse secondary antibody was used to detect the protein bands on Odyssey Clx imager (LICOR Biosciences Inc., Lincoln, NE). The protein levels were quantified using Image Studio Lite software (LICOR Biosciences Inc., Lincoln, NE).

**Cyclic-di-GMP quantification using flow cytometry.** Strains harboring the c-di-GMP-dependent $P_{CdrA}$-*gfp* transcriptional reporter plasmid subcultured into K10T-1 media and incubated at 30ºC for 24 hours. Strains without the plasmid were used as a non-fluorescent control to account for any background fluorescence. Cells were then washed, diluted, and analyzed on a Beckman 534 Coulter Cytoflex S as described previously in (8). FlowJo software version 10.8.1 was used to gate on populations of single cells. The GFP fluorescence from the $P_{cdrA}$ promoter was then measured on the gated population. A workflow of the gating strategy is described in (8).

**Supplementary Table 1**: **Strains and plasmids used in this study.**

| Strains | SMC Strain # | Description | Reference |
|---|---|---|---|
| ***Escherichia coli*** | | | |
| S17-1(λ-pir) | SMC117 | *thi pro hsdR*- *hsdM*+ ∆*recA* RP4-2::TcMu-Km::Tn7 | (9) |
| ***Pseudomonas fluorescens* Pf0-1** | | | |
| ∆*lapG*∆*lapD* | SMC 6422 | Pf0-1 with unmarked deletion of *lapG* and *lapD* | (10) |
| ∆*lapG*∆*lapD lapA-HA* | SMC 5116 | ∆*lapG*∆*lapD* with internal 3x HA epitope tag on *lapA* | (10) |
| ∆*lapG*∆*lapD dvhA-RM-swap-HA* | SMC 8159 | ∆*lapG*∆*lapD lapA-HA* with N-terminal of *lapA* (aa 1-125) swapped with N-terminal of *dvhA* (aa 1-119) at the native locus. | This study |
| ∆*lapG*∆*lapD dvhA-hlx-swap-HA* | SMC 8176 | ∆*lapG*∆*lapD lapA-HA* with the known *lapG* proteolysis residues of *lapA* (aa 81-123) swapped with predicted *dvhG* proteolysis residues of *dvhA* (aa 87-119) at the native locus. | This study |
| ∆*lapG*∆*lapD dvhA-hlx-swap-HA* AA-RR | SMC 9825 | ∆*lapG*∆*lapD dvhA-hlx-swap-HA* with 105PAAG108 mutated to 105PRRG. | This study |
| ∆*lapG*∆*lapD dvhA-hlx-swap-HA lapB*::pMQ89 | SMC 9826 | ∆*lapG*∆*lapD dvhA-hlx-swap-HA* with *lapB* gene disrupted on the *P. fluorescens* chromosome. | This study |
| ∆*lapG*∆*lapD dvhA-hlx-swap-HA* attTn7:: *lacI*-$P_{tac}$*dvhD-HA* | SMC 9827 | ∆*lapG*∆*lapD dvhA-hlx-swap-HA* with *lacI* and *dvhD-HA* under $P_{tac}$ promoter inserted at the *P. fluorescens* attTn7 site. | This study |
| ∆*lapG*∆*lapD*∆*rapA dvhA-hlx-swap-HA* attTn7:: *lacI*-$P_{tac}$*dvhD-HA* | SMC 9828 | ∆*lapG*∆*lapD dvhA-hlx-swap-HA* attTn7:: *lacI*-$P_{tac}$*dvhD-HA* with unmarked *rapA* deletion. | This study |
| ∆*lapG*∆*lapD* $P_{xutR}$-rapA *dvhA-hlx-* | SMC 9829 | ∆*lapG*∆*lapD dvhA-hlx-swap-HA* attTn7:: *lacI*-$P_{tac}$*dvhD-HA* with rapA under the control of xylose-inducible promoter $P_{xutR}$ at its native locus | This study |

| | | | |
|---|---|---|---|
| *swap-HA* attTn7:: *lacI*-$P_{tac}$*dvhD-HA* | | | |
| **Plasmids** | | **Description** | **Reference** |
| pMQ30 | SMC 2765 | Vector for cloning in yeast and allelic exchange in Gram-negatives; Gentamycin-resistance ($Gm^r$) cassette | (11) |
| pMQ30-*dvhA-RM-swap* | SMC 8158 | Vector to replace aa 1- 125 of *lapA* with aa 1-119 of dvhA using allelic exchange; $Gm^r$ | This study |
| pMQ30-*dvhA-hlx-swap* | SMC 8175 | Vector to replace aa 81-123 of *lapA* with aa 87-119 of *dvhA* using allelic exchange; $Gm^r$ | This study |
| pMQ30-*dvhA-hlx-swap* AA-RR | SMC 9831 | Vector to replace 105PAAG108 with 105PRRG108 in *dvhA-hlx-swap* using allelic exchange; $Gm^r$ | This study |
| pMQ56-mTn7 | SMC 3242 | Shuttle vector for insertion of the mini-Tn7 element from pUT18-mTn7-Gm into the attTn7 site on the *P. fluorescens* chromosome; Carbenicillin-resistance ($Cb^r$) cassette | (11) |
| pBUX-BF13 | SMC 1822 | Helper plasmid encoding Tn7 transposition functions; $Cb^r$ | (12) |
| pFLP3 | SMC 4060 | Vector for removal of antibiotic resistance cassettes flanked by FRT sites with flipase; *sacB* for counterselection; $Cb^r$ | |
| pMQ56-*lacI*$P_{tac}$-*dvhD-HA* | SMC 9832 | Vector for integration of *lacI* repressor and HA-tagged *dvhD* under IPTG-inducible Ptac promoter at the attTn7 site on the P. fluorescens chromosome; $Cb^r$ | This study |
| pMQ30-*rapA* | SMC 9833 | Vector to produce an unmarked deletion of *rapA* from *P. fluorescens* chromosome by allelic exchange; $Gm^r$ | This study |
| pMQ72 | SMC 2795 | Expression vector for cloning in yeast, *E. coli* and *P. fluorescens* with arabinose-inducible gene expression system; $Gm^r$ | (11) |
| pMQ72-*dvhG-His* | SMC 9834 | Vector for arabinose-inducible expression of full length *dvhG* with a 6x-His tag at the N-terminal; $Gm^r$ | This study |
| pMQ72-*dvhG*Δ*TM-His* | SMC 9835 | Vector for arabinose-inducible expression of *dvhG* without aa 1-40 and with a 6x-His tag at the N-terminal; $Gm^r$ | This study |

| | | | |
|---|---|---|---|
| pMQ72-*dvhG-His C317A* | SMC 9836 | Vector for arabinose-inducible expression of full length *dvhG-C317A* with a 6x-His tag at the N-terminal; $Gm^r$ | This study |
| pMQ72-*lapG* | SMC 5143 | Vector for arabinose-inducible expression of full length *lapG*; $Gm^r$ | (10) |
| pMQ89 | SMC 2952 | Suicide vector to generate single-crossover constructs in *P. fluorescens* chromosome; $Gm^r$ | (11) |
| pMQ89-*lapB* | SMC 5148 | Vector to introduce disruption in *lapB* gene on *P. fluorescens* chromosome; $Gm^r$ | (13) |
| pMQ650 | SMC 9396 | Expression vector for cloning with xylose-inducible gene expression system; $Gm^r$ | (14) |
| pCdrA-gfpC | SMC 5709 | Vector for fluorescence-based quantification of cyclic di-GMP in *Pseudomonas aeruginosa* and related species | (15) |

**Supplementary Table 2: Oligonucleotides used in this study.**

| Oligonucleotide Name | Sequence (5' → 3') |
|---|---|
| DvhA_RMswap_up_F | GTCGACTCTAGAGGATCCCCCGCTGTCGGTGCCTTCAC |
| DvhA_RMswap_up_R | GATGGTGCGATTGAGAGGCATTGAAGACTCTCCGGGTGTCAC |
| DvhA_RMswap_down_F | GCAGGTAGCGGCGGTGCCGCTGGTGGCGGC |
| DvhA_RMswap_down_R | CGAATTCGAGCTCGGTACCCCCTTCGGTCACGCTTGG |
| DvhA_RMinsert_F | GTGACACCCGGAGAGTCTTCAATGCCTCTCAATCGCACCATC |
| DvhA_RMinsert_R | GTCGCATCGAGCATCACGAAACCGCCGCTACCTGC |
| DvhA_hlxswap_up_F | GTCGACTCTAGAGGATCCCCCGCTGTCGGTGCCTTCAC |
| DvhA_hlxswap_up_R | CCGGGGAGTTCCTGACCGGTGGCTTCGGCCAGGTC |
| DvhA_hlxswap_down_F | GCAGGTAGCGGCGGT ACTGGCGGTGCCGCT |
| DvhA_hlxswap_down_R | CGAATTCGAGCTCGGTACCCCCTTCGGTCACGCTTGG |
| DvhA_hlxInsert_F | GACCTGGCCGAAGCCACCGGTCAGGAACTCCCCGG |
| DvhA_hlxInsert_R | CCAGCGGCACCGCCACCGCCGCTACCTGC |
| DvhG_nHis _F | TCGAGCTCGGTACCCGAAGGAGATATACATATGCATCATCACCATCAC CACGCAAAGGCGCGGGAGA |
| DvhG _R | GACTCTAGAGGATCCCCTCATTTCCTCCCGGCTTTCTTC |
| DvhG-noTM_nHis _F | TCGAGCTCGGTACCCGAAGGAGATATACATATGCATCATCACCATCAC CACGGAGATACCGAAGCTGGGG |
| delrapA_up_F | GTCGACTCTAGAGGATCCCCCCGGCGAGCGATCCGTTC |
| delrapA_up_R | GTTTCGTGCATTTGAAGAGGGGCAATCTCTGGCGATAAAAAAAGG |
| delrapA_down_F | ATCGCCAGAGATTGCCCCTCTTCAAATGCACGAAACGGC |
| delrapA_down_R | CGAATTCGAGCTCGGTACCCGGGCGGAAGTGAAATTTGATAAACG |
| rapAup_F | GTCGACTCTAGAGGATCCCCCACGTAGTCCGGCCGC |
| rapAup_R | CCAAGAACAACAAGGAGGATTTTATGACCACGACCGAACAG |
| PxutR_insert_F | GCTGTTCGGTCGTGGTCATAAAATCCTCCTTGTTGTTCTTGG |
| PxutR_insert _R | ATCGCCAGAGATTGCCCGAGCCCTATCGGCTGG |
| rapAdown_F | CCAGCCGATAGGGCTCGGGCAATCTCTGGCGATAAAAAAAGG |
| rapAdown_R | CGAATTCGAGCTCGGTACCCCGGCGAGCGATCCGTTC |
| LapB_seqF | GGCCGAGATCAAAAACATTCAGTTAT |
| LapB_seqR | TGGCCCACAGCATCAGG |
| Fra1_PtaclacI-dvhD_mTn7_F | CTCACTAGTGGATCCCCCTCACTGCCCGCTTTCCAGTC |
| Fra1_PtaclacI-dvhD _mTn7_R | ACAGGAAACAGACTAGTGCTCTGCAGAAGGAGATATACATATGTACCCATAC |

| | |
|---|---|
| Fra2_PtaclacI-dvhD _mTn7_F | ACAGGAAACAGACTAGTGCTCTGCAGAAGGAGATATACATATGTACCCATAC |
| Fra2_PtaclacI-dvhD _mTn7_R | GGAATTCCTGCAGCCCTCAGCCTTGACGCACATCAAGC |
| hlxswap_RR_up_F | GTCGACTCTAGAGGATCCCCGGCAACCGGAATGACCGTATG |
| hlxswap_RR_up_R | CGGCGGTCGCATCGAGCAT |
| hlxswap_RR_down_F | ATGCTCGATGCGACCGCCG |
| hlxswap_RR_down_R | CGAATTCGAGCTCGGTACCCGATGGTGTCGGTGACCTGAGTC |
